# Dynamic modulation of genomic enhancer elements in the suprachiasmatic nucleus mediates daily timekeeping in mammals

**DOI:** 10.1101/2022.11.15.515402

**Authors:** Akanksha Bafna, Gareth Banks, Michael H. Hastings, Patrick M. Nolan

## Abstract

The mammalian suprachiasmatic nucleus (SCN), located in the ventral hypothalamus, is crucial for synchronising and resetting all cellular rhythms in accordance with critical environmental and visceral cues. Consequently, the systematic regulation of spatiotemporal gene transcription in the SCN is vital for daily timekeeping. Here, we sought to identify SCN enriched gene regulatory elements that enable temporal gene expression using histone-ChIP-seq. We found a vast majority of SCN enhancers not only exhibit robust 24-hour rhythmic modulation in H3K27ac occupancy, but also possess canonical E-box (CACGTG) motif, potentially influencing downstream cycling gene expression. In parallel, we conducted RNA-Seq at six distinct times to establish enhancer-gene relationships in the SCN. Surprisingly, around 35% of cycling H3K27ac abundance is seen adjacent to rhythmic gene transcripts, often preceding the rise in mRNA levels. We also noted that enhancers encompass non-coding actively transcribing enhancer RNAs (eRNAs), that in turn oscillate along with cyclic histone acetylation to direct gene transcription. Taken together, these findings shed light on genome-wide pre-transcriptional regulation operative in the central clock that enables its orchestration of daily timekeeping mechanisms in mammals.

## Introduction

Most organisms possess intrinsic circadian (approximately one day) clocks that align their molecular, behavioural and physiological processes to changing daily environmental conditions. In mammals, the phase and amplitude of circadian processes are directed and synchronized by the suprachiasmatic nucleus (SCN) of the hypothalamus, which facilitates robust 24-hour oscillations in peripheral tissues (Hastings et al., 2018; Schibler et al., 2015). In turn, the SCN receives retinal input that ensures its autonomous circadian time is synchronised to solar time. Circadian time is generated by a genetic network of cell-autonomous transcription-translation loops (TTFLs) driven by the activators CLOCK (circadian locomotor output cycles kaput) and BMAL1 (brain and muscle ARNT-Like 1) and repressors PER (period) and CRY (cryptochrome) (Takahashi et al., 2008). At the circuit-level, intercellular synchrony in the SCN is maintained through neuropeptidergic systems such as vasoactive intestinal peptide (VIP) and arginine vasopressin (AVP), which serve as coupling factors (Herzog et al., 2017). Ablation of the SCN leads to perpetual loss of circadian timing with out-of-sync peripheral clocks (Rusak & Zucker, 1979; Yoo et al., 2004), whereas genetic restoration of the SCN TTFL has been shown to initiate 24 hour rhythmicity at the molecular and behavioural levels (Maywood et al., 2021).

Precise regulation of spatiotemporal gene expression in the SCN is imperative for daily timekeeping. Daily rhythms of electrical activity, metabolic rate and intra- and inter-cellular signalling are driven by cascades of transcription, co-ordinated by clock controlled genes (CCGs) many of which are transcription factors sensitive to the TTFL. Such rhythmic gene transcription depends on the sequential co-ordination of multiple layers of epigenetic events ranging from chromatin accessibility to transcription factor (TF) binding (Yeung et al., 2018). Interestingly, enhancer-promoter interactions have been shown to govern RNA polymerase II (RNAP II) recruitment followed by TF and co-factor binding, leading to cell- and tissue-specific expression through the initiation of mRNA transcription. In mouse liver, for example, 24-hour periodic occupancy by chromatin remodelling enzymes permits RNAP II and BMAL1 recruitment to support circadian gene expression (Koike et al., 2012; Sobel et al., 2017). Additionally, advances in genome sequencing technologies have highlighted the occurrence of rhythmic modulation in chromatin topology that facilitate proximal and distal circadian gene expression (Aguilar-Arnal et al., 2013; Kim et al., 2018; Mermet et al., 2018) through promoter-enhancer “looping”.

Recently, active enhancer sites have also been reported to transcribe enhancer RNA (eRNA) dynamically in response to chemical and electrical stimuli, learning and behavioural experiences (Joo et al., 2016; Kim et al., 2010; Telese et al., 2015). This class of non-coding RNAs are generally not spliced or polyadenylated (Arner et al., 2015; Gray et al., 2015) and is believed to assist long distance enhancer-promoter interaction. Incidences of circadian eRNAs oscillating and peaking with distinct phases have been observed in the mouse liver transcriptome (Fang et al., 2014). Such circadian variation in abundance of eRNA adds an important layer of control in the regulation of gene expression in defined time and space.

At the tissue level, the persistence and precision of the SCN as a circadian time-keeper are unique, but the cell-autonomous regulatory factors that confer this uniqueness to the SCN are poorly understood. Genomic mapping of regions with associated histone modifications such as H3K4me3 and H3K27ac have been the cornerstone for identifying active promoter and enhancer sites, respectively (Heintzman et al., 2007; Won et al., 2008). Typically, during active transcription, looser chromatin structures facilitate binding of transcription factors and RNA polymerases to trigger target gene expression. For instance, acetylation of a lysine residue of H3 histone weakens the binding between the histone and negatively charged DNA and exposes the DNA to regulatory proteins (Dong & Weng, 2013; Starks et al., 2019). Trimethylation of the lysine at the fourth position of the H3 histone is another conspicuous modification that is seen largely near transcription start sites (TSS) of the transcribing genes, marking the promoter regions.

Here, we used histone chromatin immunoprecipitation sequencing (Histone ChIP-seq) to map genome-wide gene promoter and active enhancer sites in the SCN. Next, with a view to identifying putative SCN-enriched regulatory processes, we compared the identified histone modification peak profiles in the SCN with those found in cerebral cortex. This revealed considerable enrichment associated with genes highly expressed in SCN (Brown et al., 2017) supporting the view that histone modifications are reliable markers for mapping gene regulatory elements in a locus-, cell- and tissue-specific manner (Garcia et al., 2008; Ngo et al., 2019). We then turned our attention to the 24-hour oscillatory patterns at the identified gene regulatory *cis*-elements, which presumably are vital to drive the SCN circadian rhythm. Differential histone modification analysis across day and night revealed robust periodic occupancy of H3K27ac, thereby marking cycling active enhancer sites in the SCN. In contrast, the promoter counterpart, H3K4me3 occupancy, was not as dynamic. Furthermore, we carried out directional bulk RNA-seq at six distinct time-points to study the relationship between observed rhythmic H3K27ac occupancy and target gene expression. Along with rhythms in the coding mRNA fraction, we also noticed cycling expression in the non-coding bidirectional enhancer RNA (eRNA). This was related to differential H3K27ac sites that can potentially influence both proximal and distal circadian gene expression (Li et al., 2016; Sanyal et al., 2012).

Our findings offer a framework to study the intricate dynamic pre-transcriptional regulation operative in the central clock that will contribute to maintaining its daily rhythms. The marked modulation in H3K27ac occupancy exposes enhancer sites, and sets a wave of chromatin accessibility indispensable for subsequent TF and co-factor binding along with RNAP II. This timely availability of a transcriptional apparatus enables the pace of rhythmic gene transcriptional machinery in the SCN to be set. To our knowledge, this is the first study that highlights SCN-enriched enhancers and demonstrates periodic variation in such chromatin modifiers linked to circadian gene expression.

## Results

### Genome-wide promoter and enhancer site mapping in the SCN using histone ChIP-Seq

To identify DNA regulatory elements influencing gene transcription in the SCN, we focussed on SCN-enriched promoter and enhancer sites marked by H3K4me3 and H3K27ac, respectively. C57BL/6J mice were kept in standard 12hr light-12hr dark conditions and eight animals per time-point (Material and methods) were used for subsequent brain dissections. The harvested SCN and cerebral cortical brain regions were then used for H3K4me3 and H3K27ac ChIP-Seq and the resulting sequencing reads were mapped to the mouse genome (Materials and methods). Initial principal component analysis (PCA) of aligned reads arising from H3K4me3 and H3K27ac occupancy in duplicate biological samples confirmed high quality ChIP signal and tissue-specific enrichment. Subsets of reads associated with H3K4me3 and H3K27ac were not only found to be separate from their corresponding input sample, but also showed clear distinction based on the brain region (Fig 1A). In addition, the variability between tissue-specific (SCN vs. cortex) H3K27ac marks was greater than the H3K4me3 counterpart, suggesting a potential role for active enhancers marked by H3K27ac in defining tissue identity and function (Ko et al., 2017). As expected, the occupancy of H3K4me3 (along with H3K27ac) was highly abundant around TSSs (TSS ± 2Kb) presumably arising from the actively transcribing genes (Fig S1) (Beacon et al., 2021). A collection of 10,577 H3K4me3 peaks (FDR≤0.05) was observed to differ in abundance significantly between SCN and cortex, with the majority located in the promoter region (Fig S2A, B, Table S1). Unsurprisingly, SCN-enriched H3K4me3 sites were found adjacent to TSS of the genes known to be highly expressed in the tissue, as exemplified in Fig. S2C (Brown et al., 2017; Wilcox et al., 2017), while many were associated with synaptic processes and signalling pathways (Fig S2D).

**Fig. 1.**
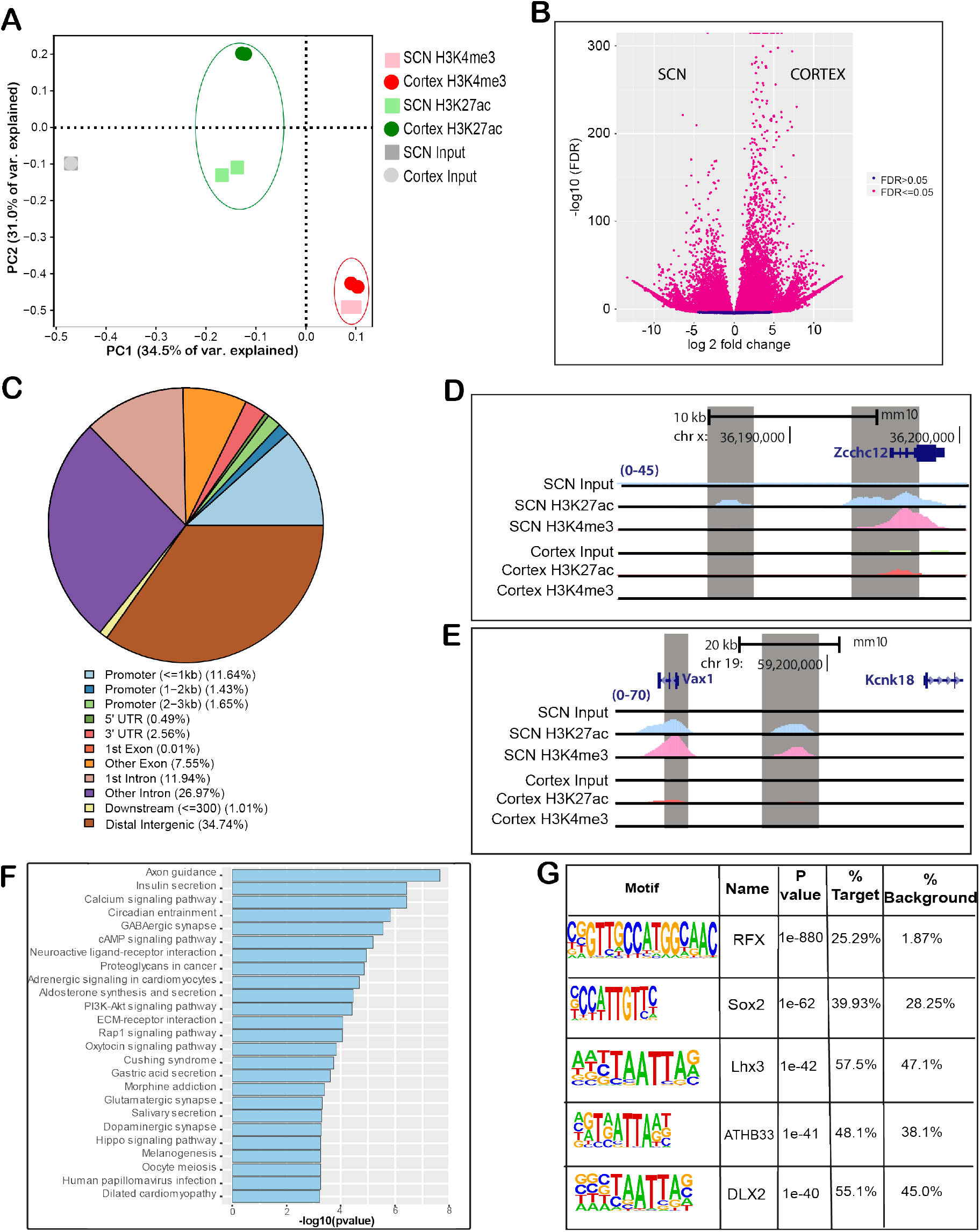
Genome-wide characterization of SCN enhancers. (A) PCA analysis of aligned reads post H3K4me3 and H3K27ac immunoprecipitation from SCN and Cortex mouse brain tissues. (B) Volcano plot showing fold change and false discovery rate (FDR) for differential H3K27ac sites between SCN and Cortex as computed by Diffbind. (C) Genomic feature distribution of SCN enriched H3K27ac peaks (n= 14153). (D,E) UCSC genome browser tracks showing histone modifications H3K4me3 and H3K27ac normalized ChIP-seq read coverage (shaded grey) along with their input for SCN and Cortex at representative examples. The chromosome location and scale (mm10 genome) indicated at the top. (F) Functional annotation of nearest neighbouring gene (TSS) to SCN enriched H3K27ac sites (fold change > 5) using KEGG pathway (DAVID). (G) Over-represented transcription factor binding motifs using SCN enriched H3K27ac sites as target by HOMER.

Having identified SCN-specific promoter marks via H3K4me3, we then sought to delineate active enhancers by mapping H3K27ac binding. We identified approximately four times more H3K27ac peaks (44,247 sites, FDR≤0.05) that differentially marked active enhancer sites between the two brain regions (Fig 1B, Table S2). Interestingly, SCN-enriched H3K27ac sites were dominant at distal intergenic (34.7%) and intronic regions including first introns (38.9%) (Fig 1C), which is consistent with previous reports (Creyghton et al., 2010; Rada-Iglesias et al., 2011) wherein tissue-specific enhancer marks were seen to be highly abundant at intergenic and intragenic regions. Next, we preferentially selected highly distinct (fold change ≥ 5, Fig. 1D, E) SCN-specific H3K27ac sites and recorded the closest TSS, guided by peak annotation tool ChIPseeker (Material and methods). Notably, many of the genes observed to be adjacent to these active enhancer marks were found to be implicated in SCN-enriched functions, including circadian entrainment, calcium signalling and GABAergic synaptic function (Fig 1F) (Koronowski & Sassone-Corsi, 2021; Morris et al., 2021) Furthermore, motif analysis on H3K27ac-bound regions (Material and Methods) revealed enrichment of the RFX and SOX families of transcription factor binding sites that are known to be present in adult SCN tissue and influence light entrainment pathways (Araki et al., 2004; Cheng et al., 2019) (Fig 1G) Overall, we produced a genome-wide distribution of SCN enhancer marks (Fig 2, layer 1-3) along with the neighbouring target gene (layer 4), which likely supports the role of the SCN as the principal circadian pacemaker. The SCN-enriched enhancer sites is seen to be well-distributed across the genome, with high incidence of the enriched sites (fold change > 10, layer 3 of Fig.2) at certain genomic locations such as chromosome (chr) 7 and chromosome 17. Interestingly, many of these SCN-enriched enhancers were also found clustered together, as seen within *Usp29* (± 0.2 Mb) gene at chr 7 and *Zfhx3* (± 1Mb) gene at chr 8. It is noteworthy that the SCN transcription factor *ZFHX3* have been implicated as an important regulator in setting the pace of the circadian clock (Parsons et al., 2015), and prevalence of SCN-enriched enhancer marks hints on its tissue-specific transcriptional regulation.

**Fig. 2.**
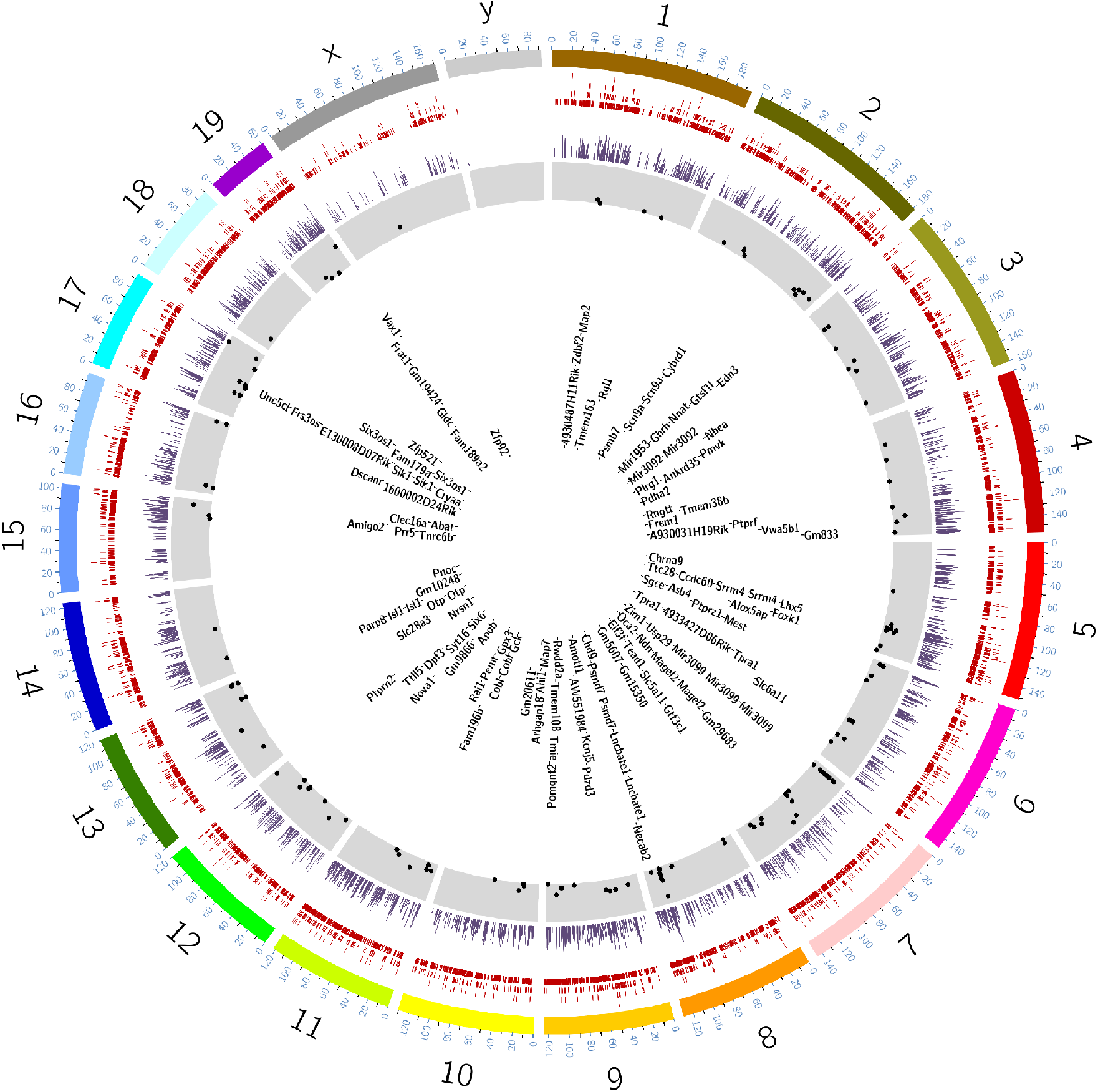
SCN Enhancer map. Circos plot for SCN enriched enhancers with mm10 genome assembly and four layers (outer to inner), layer 1: SCN enriched H3K27ac sites (n = 14153), layer 2: Histogram showing SCN enriched H3K27ac sites with fold change > 5 in comparison to Cortex, layer 3: Scatter plot showing subselected SCN enriched H3K27ac sites with fold change >10 and layer 4: Nearest gene (TSS) to the sites mapped in layer 3.

### Time-of-day dependent variation in enhancer activity in SCN

The potential regulation of the abundance of histone modifications (H3K4me3 and H3K27ac) was then investigated by comparing the anti-phasic (12-hour apart) time-points, ZT3 (3 hours after light on) and ZT15 (3 hours after light off) in both SCN and cerebral cortex. We observed approximately 551 differential H3K4me3 sites in cortex at FDR ≤ 0.05 (Fig S3A, Table S3A), mostly confined around the TSS of genes involved in learning and synapse organization. In contrast, no changes in H3K4me3 sites were seen within the SCN. This stability of histone trimethylation levels in the SCN is in marked contrast to studies in peripheral clock(s) (Koike et al., 2012; Le Martelot et al., 2012; Valekunja et al., 2013) wherein occupancy of the promoter mark was shown to be cyclic. Rhythmic trimethylation may therefore be a conserved feature of driven clocks in the cortex and the peripheral tissues. In contrast, there was strong differential H3K27ac occupancy within the SCN between day (ZT3) and night (ZT15) (Fig. 3A, Table S4A). This reinforces the crucial role of H3K27ac marked active enhancers, as opposed to H3K4me3 marks, in regulating time-of-day dependent gene transcription. On further inspection, a significant proportion of identified differentially H3K27ac marked enhancer sites were found proximal to genes implicated in circadian entrainment, such as *Gnas, Rasd1, Creb1* (Brown et al., 2017; Lee et al., 2010) (Table S3B,C Fig. 3B). Therefore, we could precisely identify time-of-day dependent H3K27ac variance in the SCN, as a potential contributor to, and/ or output from, daily TTFL timekeeping. We then examined whether the observed differential H3K27ac sites in SCN are also present in cortex, even though they may not be time-dependent (ZT3 vs. ZT15). Notably, about half of these differential sites (n= 146) revealed greater H3K27ac abundance in SCN (Fig. 3C, S3B) with almost negligible occurrence in cortex. Moreover, active enhancer sites that showed day-night fluctuation in both SCN and cortex (n =31) did not follow the same trend, for example a differential H3K27ac site mapped to calcium binding gene *Calm3* is strongly elevated at ZT3 in the SCN, but is elevated at ZT15 in the cortex (Fig. 3D), where it may potentially control opposing peaks of gene expression (Chun et al., 2015). Hence, this flux in H3K27ac occupancy suggests a role of *cis*-regulatory elements in directing the tissue-specific gene transcriptional machinery in response to environmental and cell-autonomous stimuli. It also highlights the possibility of same enhancer site being active at distinct time-of-day to regulate the subsequent tissue-specific gene expression.

**Fig. 3.**
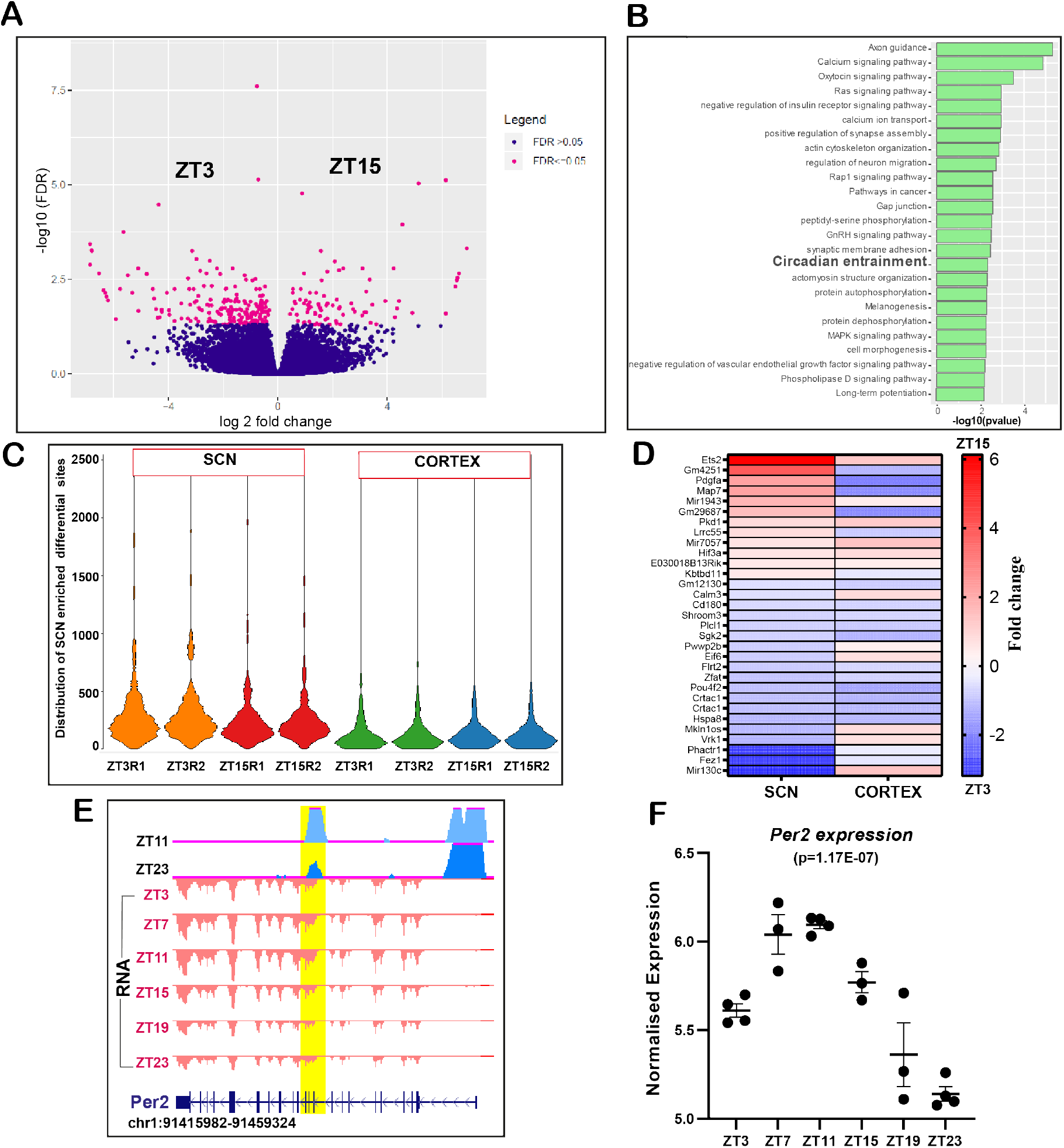
Dynamic occupancy of H3K27ac based on time of the day. (A)Volcano plot showing fold change and false discovery rate (FDR) for differential H3K27ac sites in SCN between ZT3 and ZT15 (n = 293) as computed by Diffbind. (B) Functional annotation of nearest gene (TSS)-from differential (ZT3 Vs ZT15) H3K27ac sites using GO (biological process) and KEGG pathway (DAVID). (C) Violin plot showing distribution of differential H3K27ac peaks at ZT3 and ZT15 in SCN and cortex. (D) Heat map view of differential H3K27ac sites (ZT3 Vs ZT15) observed in both SCN and cortex. (E) UCSC genome browser tracks showing differential (fold change =1.7) H3K27ac intragenic peak (blue with shaded yellow) at the *Per2* gene locus between ZT11 and ZT23 along with the gene expression tracks (red) across six time points (ZT3-ZT23) arising from the negative strand. (F) Normalised expression level of the *Per2* gene across six distinct time-points, cycling with a peak at ZT11 and a trough at ZT23, Pearson correlation (H3K27ac peak and gene expression) = 1.

To understand further the dynamics of H3K27ac occupancy in the context of daily timekeeping in the SCN, we profiled its occurrence every four hours starting from ZT3 (Material and Methods) and compared the peak profiles between further anti-phasic time-points (ZT7 Vs ZT19, ZT11 Vs ZT23), based on the expectation that rhythmic genes under circadian regulation would have peaks and troughs in expression level approximately 12 hours apart (Fig. S3C). Indeed, we found a considerable number of active enhancer sites in the SCN (Table S4B, C) with differential abundances at distinct times of day. Surprisingly, there was almost negligible overlap between H3K27ac marked differential sites noticed from distinct anti-phasic comparisons, for instance an enhancer peak differential between ZT3 and ZT15 did not overlap with any found between ZT7 and ZT19 i.e., events were temporally segregated. Using this approach, we clearly identified a unique active enhancer site within the *Per2* gene showing increased histone acetylation at ZT11 in comparison with ZT23, coinciding with the peak and trough of host gene expression levels (Fig. 3E, F). Similarly, a differential H3K27ac peak (ZT7 Vs ZT19) observed upstream of the *Cry1* TSS is concordant with the downstream gene transcription (Table S4B). Thus, variability in histone acetylation levels seems to be clearly associated with changing transcriptional output in the SCN, as noted with rhythmic expression of core-clock genes; *Per2 and Cry1*. About 50% of the differing histone acetylation levels perfectly correlated (Pearson correlation =1) with the host gene expression (Fig. S4E). To our knowledge, this method of strategic in-depth examination in the time-dependent occurrence of histone modifications has never been reported previously. We clearly observed the time-dependent differing histone acetylation levels from arising from TTFL components to downstream CCGs in the SCN.

Motif analysis of anti-phasic differential H3K27ac sites revealed the pre-dominance of distinctive transcription factors as a function of the time of the day (Fig. S3D). Transcription factor binding sites for CAAT/enhancer-binding protein (C/EBP) and retinoic orphan receptor (ROR), both belonging to the PARbZip family (Gachon, 2007), were abundant in the ZT3 vs. ZT15 comparison, whereas those for NF-kB, implicated in the mammalian circadian clock, were enriched in the ZT11 vs. ZT23 comparison (Shen et al., 2021). This is consistent with finding in the liver where phase-separated functional enhancers were driven by distinct TFs. (Fang et al., 2014). Therefore, we assume that the change in histone acetylation is important for the timely binding of TFs to drive temporal gene expression as noted across various tissues.

### Diurnal variation in H3K27ac occupancy linked to rhythmic gene transcription

To determine the prevalence of 24-hour oscillation in H3K27ac abundance, we considered peaks that showed differential occupancy between any anti-phasic (12 hours-apart) comparisons. Active enhancer sites that were found to be distinct between these were compiled (n = 1,021) and further assessed for circadian oscillation using the Extended Circadian Harmonic Oscillator (ECHO) application (De Los Santos et al., 2020). Briefly, logarithmic normalised counts per million (CPM) per peak interval was used across six distinct time-points to determine 24-hour oscillation (Material and Methods). Of these, approximately half were recorded as rhythmic, and almost a quarter (n = 286) also showed robust oscillation (p ≤ 0.05) by both JTK_CYCLE (Hughes et al., 2010) and ECHO (Fig. 4A, S4A, Table S5). Interestingly, the canonical E-box *(CACGTG)* element facilitating circadian transcriptional activation by binding of CLOCK and BMAL1 [41] was over-represented among the periodic H3K27ac sites (Fig. 4C). In principle, acetylation of the lysine residues of histones is known to promote binding of transcription factors by weakening histone-DNA interactions [42]. Thus, the occurrence of rhythmic histone acetylation potentially triggers a wave of chromatin accessibility and facilitates binding of clock TFs (E-box) to regulate cyclic gene transcription. Indeed, we noticed approximately 35% of genes proximal to rhythmic H3K27ac sites were strongly cyclic (JTK_CYCLE, p < 0.05), suggesting a pivotal role of *cis*-regulatory elements in the maintenance of rhythmic transcriptional output in the SCN (Fig. 4D and F, Table S6). For example, *Rora* (RAR-related orphan receptor alpha) is known to regulate the rhythmic expression of *Bmal1* (Akashi & Takumi, 2005), and it peaks following the periodic rise of nearby histone acetylation levels as seen in Fig. 4F. This timely coordination in gene transcription provided by the adjacent histone modifiers highlights the importance of dynamic pre-transcriptional regulation in the SCN. Considering the cycling gene expression in the SCN peaks at varied times during the day (Fig. S4B, Table S7), we also observed uniform phase (peak abundance) distribution for the rhythmic H3K27ac sites (Fig. 4B). Although a significant proportion of these oscillating sites were found at intragenic regions, they were well separated from the corresponding promoters or TSS, demarcating them from any H3K27ac (along with H3K4me3) signal arising directly due to the host gene transcriptional activity (Fig. S4C, D). For that reason, a high proportion of these rhythmic histone acetylation sites were found ± 50 Kb from the rhythmic gene TSS. In addition, functional annotation of rhythmic genes that are under the control of the phase-separated cycling H3K27ac levels in the SCN revealed diverse roles (Fig. 4E) ranging from initiation of transcription (ZT3 Vs ZT15; green plot) to protein phosphorylation (ZT11 Vs ZT23; red plot). To summarize, the observed robust oscillations in H3K27ac levels, peaking at distinct times of day, contribute to the cluster of rhythmic gene expression involved in distinct functions in the SCN. These rhythmic genes either are involved in the canonical TTFL or include downstream clock controlled genes.

**Fig. 4.**
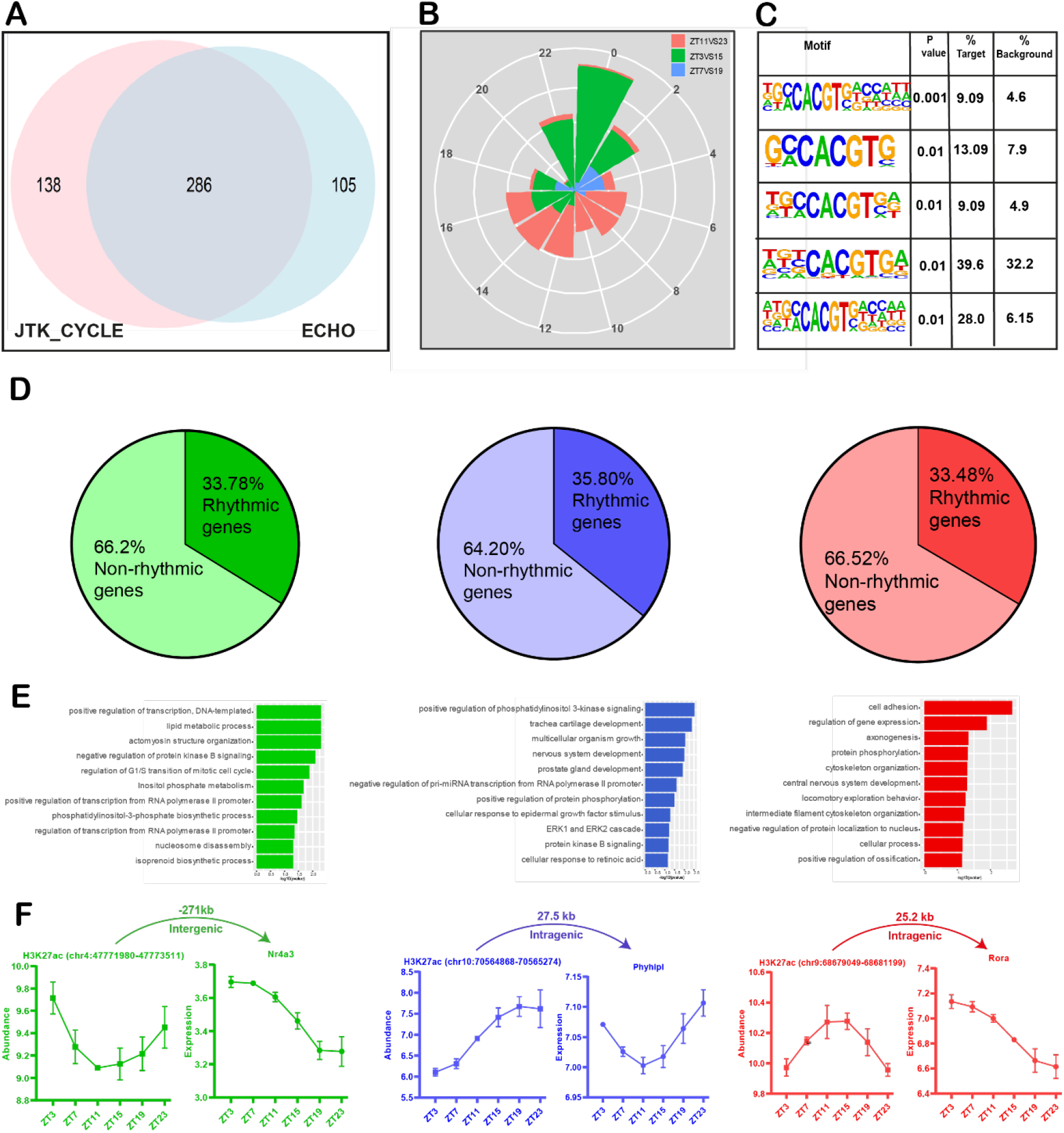
Rhythmic H3K27ac occupancy linked to cycling gene expression in SCN. (A) Venn-diagram showing overlap of rhythmic H3K27ac abundance as analyzed by ECHO and JTK_CYCLE in SCN. (B) Phase distribution of cycling H3K27ac peaks that were found differential in ZT3 Vs ZT15 (green), ZT7 Vs ZT19 (blue) and ZT11 Vs ZT23 (red) groups denoted by a rose plot. (C) Over-representation of E-box motif (CACGTG) in oscillating H3K27ac sites (HOMER). (D) Proportion of rhythmic genes mapped adjacent to rhythmic H3K27ac regions for each phase separated enhancer group. (E) Functional annotation of rhythmic genes found in close proximity of cycling H3K27ac abundance, peaking at distinct time (ZT), using GO (biological process) and KEGG pathway (DAVID). (F) Representative examples of rhythmic H3K27ac and target gene expression (p<0.05) with distance to TSS and genomic location shown at top.

### De-novo identification of actively transcribing enhancer (eRNA) in SCN

We clearly identified approximately 14,153 (FDR≤0.05) SCN-specific enhancer sites when compared to the cortex (Fig. 2). A subset of SCN enhancer marks, found at intergenic regions, were further investigated for bidirectional transcription which is considered to be the hallmark of eRNA occurrence (Fig 5A) (Material and Methods). Unlike the coding mRNA component, these long non-coding fractions (median size = 346 nucleotides) are not spliced and are not polyadenylated (Andersson et al., 2014). With the recent advancement in next generation sequencing, the occurrence of eRNA along with H3K27ac histone modification has been recognised as a reliable measure of active enhancers, and has been shown to regulate gene transcription in response to plasticity-inducing stimulation and behavioural experience (Malik et al., 2014; Telese et al., 2015). Therefore, using a direction-specific total RNA-seq SCN dataset, we revealed 2,221 intergenic bidirectional transcription sites (IBS) (Table S8). Out of these, we sub-selected IBS that revealed greater SCN-specific H3K27ac occupancy (fold change ≥ 5, HIBS) and focussed on the high confidence SCN actively transcribing enhancer sites (n=883), as represented in Fig. S5A, B. Based on previous findings, chromatin remodelling and eRNA transcription is shown to precede mRNA expression present at adjacent *cis*-loci (Arner et al., 2015; Kim et al., 2015; Schaukowitch et al., 2014). Therefore, the prevalence of eRNA in the SCN presented an excellent opportunity to investigate its influence on gene transcriptional machinery.

**Fig. 5.**
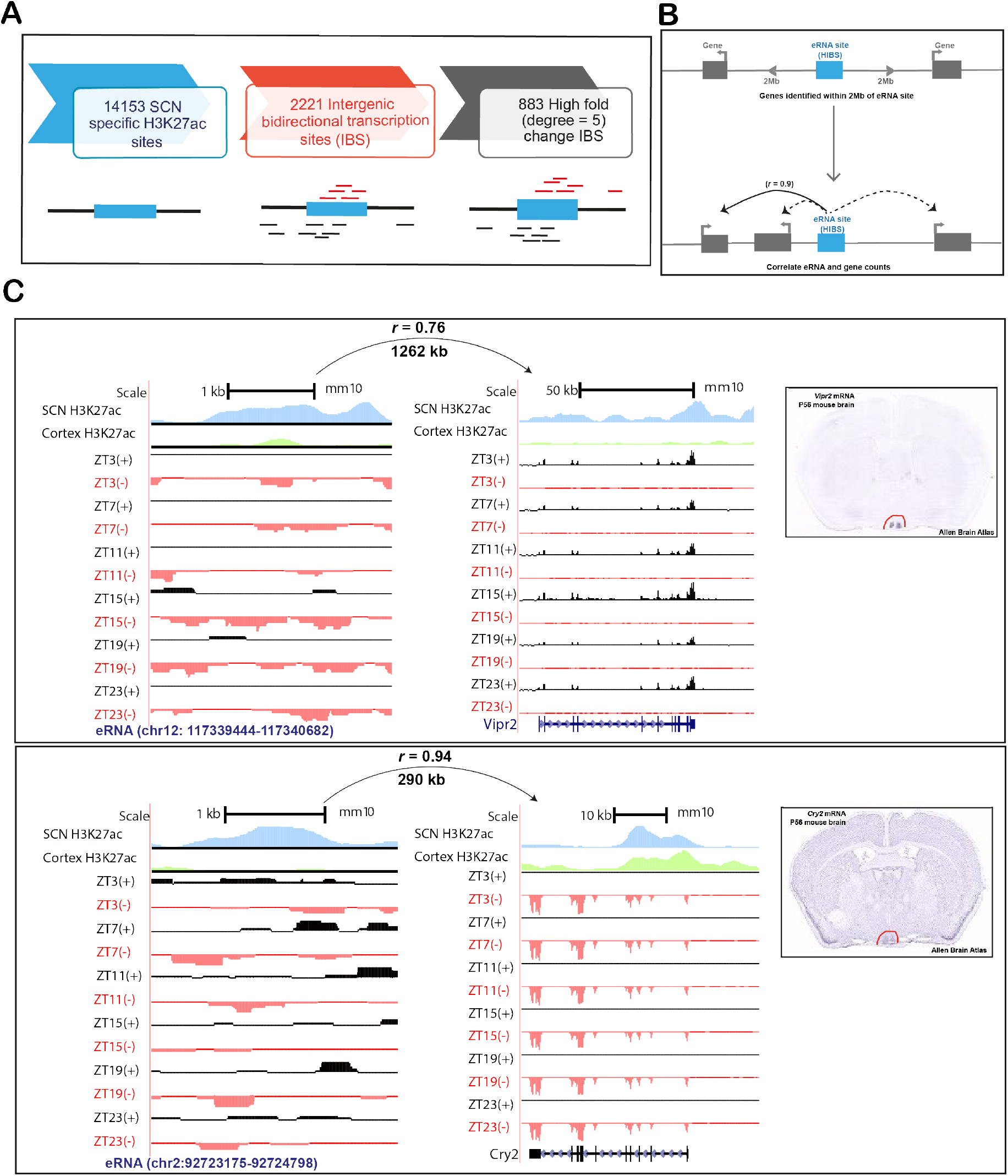
Identification of actively transcribing SCN enhancers (eRNA). (A) Analysis pipeline for identification of actively transcribing enhancers. (B) eRNA-gene pairs were determined by correlation of eRNA and gene expression levels within 2Mb distance cutoff. (C) UCSC genome browser tracks showing enhancer-linked H3K27ac signal (normalised ChIP-seq coverage) for Cortex (green) and SCN (blue) peaks and total RNA expression (reads mapped to + (black) and - (red) strand) signal as normalised CPM value for two representative examples i.e. *Vipr2, Cry2*. The distance between eRNA and predicted target gene with Pearson’s correlation is indicated above each representated eRNA-gene pair. SCN expression of the respective target gene is also evident in *in situ* hybridisation images from adult mouse brain (right, Image credit: Allen brain Atlas, Allen Institute).

Next, the expression of SCN specific eRNA was evaluated at six distinct time-points across the day starting from ZT3 (Material and Methods). In order to determine the target gene under the control of each actively transcribing enhancer, we studied the correlation between eRNA and all annotated protein-coding gene expression falling within ~±2Mb from the peak centre (Fig 5B). This approach is based on the assumption that enhancers and linked genes will correlate in their transcriptional output, as observed before across different cell classes or activity states (Carullo et al., 2020). Using positive correlation (*r* > 0.5) we noted 7,795 high confidence eRNA-gene pairs from a total of 46,022 identified pairs (Table S9). As expected, the actively transcribing enhancers positively correlated with highly expressed SCN target genes as shown in the representative examples at the *Vipr2* and *Cry2* loci (Figure 5C). Interestingly, the distance of enhancer from the target gene had no effect on subsequent strength of transcriptional correlation, consistent with the suggested role of eRNA in regulating both adjacent and distal gene expression (Han & Li, 2022) (Fig. S5C). For example, an eRNA (chr4:119682088-119683466) was found to be strongly correlated (*r* = 0.9) to both a 0.4 Mb distant target gene *Lepre1* (prolyl 3-hydroxylase 1) and a 1.5 Mb separated *Rlf* (rearranged L-myc fusion) gene. Overall, we were clearly able to show the prevalence of non-coding eRNA in the SCN, arising from enriched histone acetylation sites and well placed to regulate the tissue-specific gene expression. This finding, wherein not just the histone modifiers but also succeeding non-coding transcript could influence the target gene transcription, prompted us to study its role in daily timekeeping.

### Circadian oscillations in identified enhancer RNA expression in SCN

The identified eRNAs (Fig. 5B) from the SCN-enriched H3K27ac sites (HIBS, n = 883) were further examined for the occurrence of 24-hour oscillation in expression levels. Similar to the cycling histone acetylation abundance, we found intensely rhythmic eRNAs (Fig. 6A, JTK_CYCLE, p<0.05, Table S10) in the SCN with their relative expressions peaking across different times of the day. Based on their phase of peak expression, circadian SCN eRNAs were divided into 12 groups (phase ZT0-24, at 2 hr intervals) (Fig. 6B). To our surprise, circadian eRNAs did not cluster like the H3K27ac counterpart (Fig. 4B) or rhythmic mRNA (Fig. S4B). Approximately 66% of circadian eRNAs oscillated with a peak phase between ZT14 and ZT20, while 34% oscillated in other phases. This irregular phase distribution of SCN eRNAs agrees with the previous finding wherein circadian eRNAs in liver peaked predominantly between ZT18 and ZT3 [17]. Bearing in mind the central clock receives environmental stimuli and relays the signal to various peripheral clocks, the observed advanced phase peak in SCN circadian eRNAs relative to liver is consistent with overall phases of their respective TTFLs. By and large, the rhythmic eRNAs peaking at distinct phases, hints on the hierarchical regulation of the temporal gene expression in the SCN driven by functional enhancers.

**Fig. 6.**
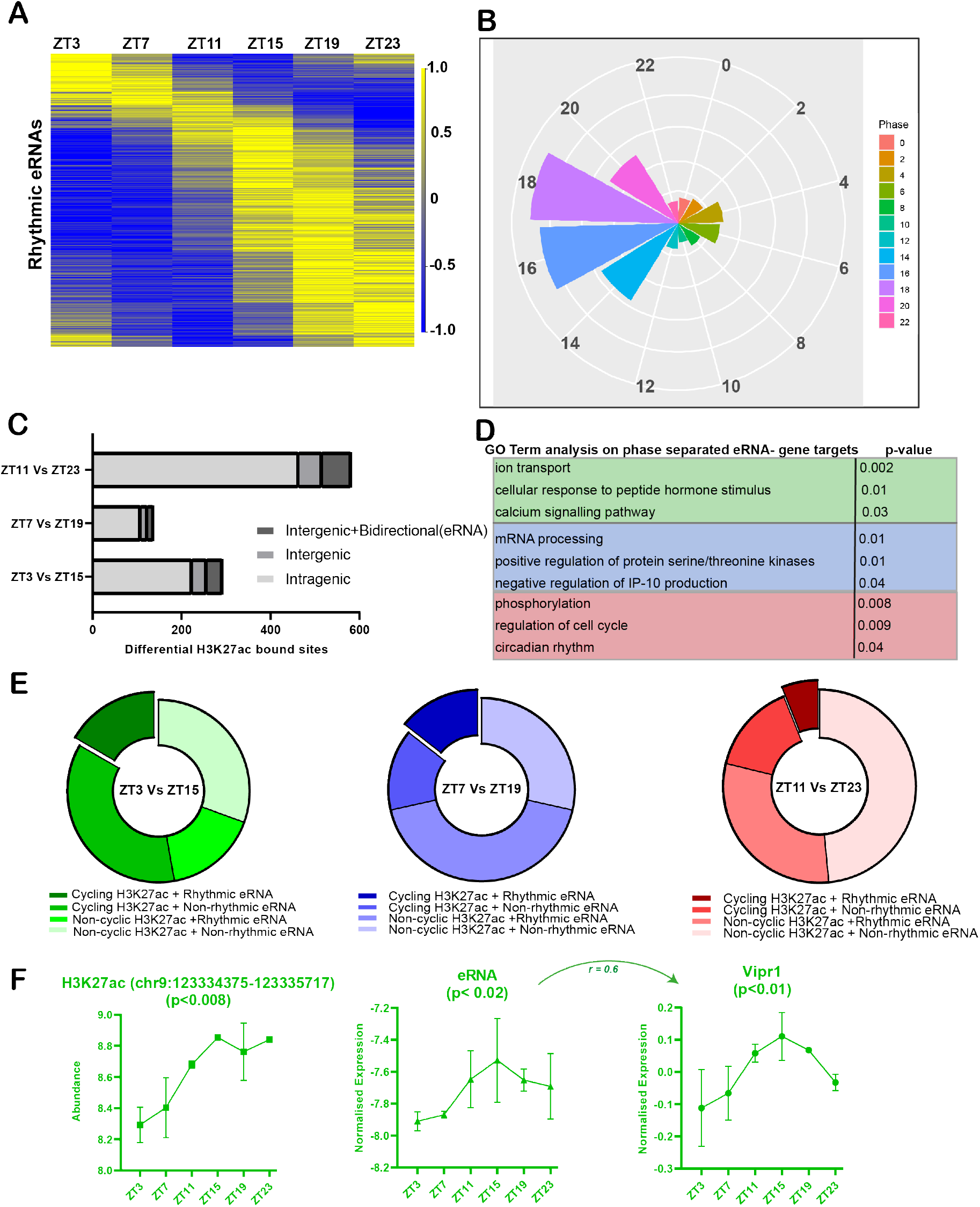
Differing H3K27ac peak abundance linked to rhythmic eRNA. (A) Heat-map showing rhythmic eRNA expression throughout the day in SCN. (B) Phase distribution of oscillating eRNA in 12 groups (2 hr interval) depicted by radial plot. (C)Proportion of differential (anti-phasic) H3K27ac sites present at intergenic regions showing bidirectional (eRNA) transcription. (D) Functional annotation of target genes controlled by eRNA arising from differential H3K27ac peak abundance at intergenic sites, ZT3Vs ZT15 (green), ZT7Vs ZT19 (red) and ZT11 Vs ZT23 (blue). (E) Distrubution of rhythmic and non-rhythmic SCN eRNAs arising from 24-hour cyclic and/ differential H3K27ac sites between compared anti-phasic group. (F) Representative example of rhythmic H3K27ac abundance, eRNA and positively correlated gene expression (log CPM) as analysed by ECHO (p<0.05) with indicated Pearson correlation coeeficient (*r*).

### Differential H3K27ac occupancy coupled with rhythmic eRNA abundance

In order to explore the link between differing histone acetylation at distinct times of day and enhancer RNA expression levels, we assessed the intergenic class of differential H3K27ac sites (n = 216) for the occurrence of eRNA transcription. To accomplish this, strand specific RNA abundance was examined for each intergenic site showing differential histone acetylation between anti-phasic time intervals. About 53% of these intergenic H3K27ac sites revealed bidirectional transcription, a reliable measure for eRNA abundance (Table S11, Fig. 6C). Subsequently, the expression level of each identified eRNA was examined across various times of the day for positive correlation (*r* > 0.5) with the target gene transcription present at ± 2Mb distance (Fig. S6A). Unlike the principal nearest neighbouring gene approach, this strategy helped us to identify a repertoire of target genes whose expression is possibly regulated by distantly transcribing enhancers in the SCN (Table S12). The recognised target genes were seen to be involved in divergent biological processes based on the disparate H3K27ac abundance as shown in Fig. 6D. For example, eRNAs identified from differential H3K27ac sites between ZT11 and ZT23 seem to regulate target genes *Arntl* (aryl hydrocarbon receptor nuclear translocator–like) (Fig. S6B), *Prokr1* (prokineticin receptor 1) etc., mediating circadian rhythms, whereas those identified from differential histone acetylation between ZT7 and ZT19 were found to influence genes involved in mRNA processing. Therefore, the non-coding eRNAs driven by temporally fluctuating histone acetylation levels adds an important layer of control over the gene expression machinery in the SCN.

Of note, about 38% of these eRNAs arising from dynamic histone acetylation sites, exhibited 24-hourly periodic transcription (ECHO/JTK_CYCLE, p < 0.05, Fig. 6E). Thus, to our knowledge this is the first time where rhythmic eRNAs were observed coupled with either cycling or differential (anti-phasic) H3K27ac peaks (Table S13). Taken together, we uncovered a proportion of intergenic sites with cycling H3K27ac levels (see layer 7, Fig. 7), which in turn transcribes rhythmic eRNA to regulate gene transcriptional machinery in the SCN. A characteristic example of this phenomenon is presented as Fig. 6F (and Fig. S6D), wherein rhythmic H3K27ac abundance and subsequent eRNA transcription is seen to regulate the circadian expression of the SCN neuropeptide receptor gene *Vipr1* (An et al., 2011). This was further confirmed by qPCR (Fig S6C) where relative expression of varying eRNA directly coordinated with the target *Vipr1* mRNA transcription during the day. As expected, the eRNA and target *Vipr1* mRNA had high expression at ZT15 and low expression at ZT3, showing an in-phase relationship. Herewith, this comprehensive analysis of dynamic histone acetylation in conjunction with enhancer transcription helped us to unfold an important layer in the systematic regulation of circadian gene expression in the SCN.

**Fig. 7.**
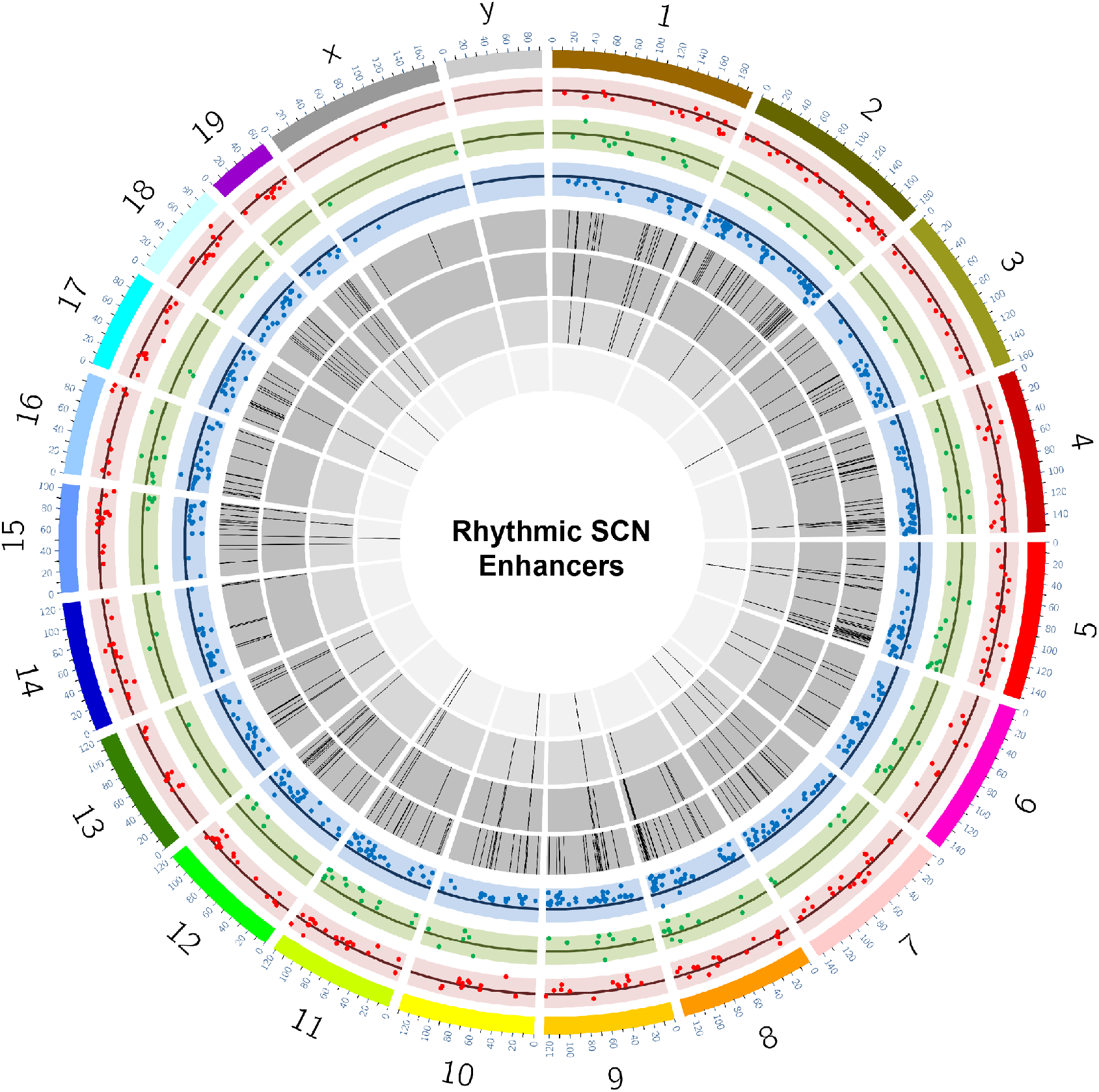
Snapshot of rhythmic H3K27ac abundance and overlapping eRNA in SCN. Circos plot for dynamic SCN enriched enhancers with mm 10 genome assembly and seven layers (outer to inner), layer 1: Differential H3K27ac sites as scatter plot (n = 293) between ZT3 and ZT15 (red), layer 2: Differential H3K27ac sites (n = 138) between ZT7 and ZT19 (green), layer 3: Differential H3K27ac sites (n = 582) between ZT11 and ZT23 (blue), layer 4: Histogram showing oscillating H3K27ac sites (n = 286, p < 0.05 both ECHO and JTK_Cycle), layer 5: H3K27ac oscillating sites from layer 4 confined to intergenic position (n = 89), layer 6: eRNA (n = 37) arising from the intergenic H3K27ac sites highlighted in layer 5, layer 7: Subset of rhythmic eRNA (n =12) co-inciding with the rhythmic H3K27ac levels mapped at the intergenic sites.

## Discussion

In mammals, circadian timekeeping is programmed by the SCN, generating daily rhythms in rest and activity, core body temperature, neuro-endocrine function, memory and psychomotor performance, and a host of other behavioural and physiological processes (Mohawk et al., 2012). The SCN is unique in its ability to sustain autonomous rhythmicity which is driven by circadian cascades of transcription that direct metabolic, electrophysiological and signalling rhythms. Our current finding highlights the instrumental role of gene regulatory elements in driving the daily timekeeping mechanism. In particular, we noted a dynamic pre-transcriptional network, operative in the SCN that potentially aids the rhythmic gene transcription and provides timely co-ordination to set the pace of the clock. First, we profiled genome-wide post-translational histone modifications to localize accessible chromatin regions that are essential for the temporal gene transcription in the SCN. With the power of the definitive H3K27ac and H3K4me3 marks, we precisely located active enhancers and gene promoters in the SCN. Furthermore, comparative assessment of the histone modifications between the SCN and cortex helped us to identify tissue-specific active enhancers and promotors and answer the longstanding question about the exclusivity of the SCN. As a result, we clearly noted abundance of tissue-specific H3K4me3 bound gene promoters (Table S1) and H3K27ac bound enhancers (Fig. 1D,E, Table S2) around TSS of the genes reported to be highly expressed in the central clock (Boguski & Jones, 2004; Brown et al., 2017; Clark et al., 2013), such as the SCN specific *Six6* gene (Fig. S2C). This led us to produce the SCN regulatory map highlighting the active enhancer sites that distinguishes SCN from the other brain region(s). Interestingly, these SCN-enriched active enhancer sites were found to be in close proximity of the genes associated with not just circadian entrainment but also calcium signalling and neuropeptidergic synaptic function (Fig. 1F); encompassing biological processes that support the daily timekeeping mechanism in the central clock. Consequently, the SCN-enriched enhancers were also seen adjacent to the genes defining SCN cell types such as neurons, astrocytes, ependymal (Wen et al., 2020; Xu et al., 2021) and holds potential to further delimit the subtypes based on the regulatory elements.

Next, we wanted to explore if any of the identified SCN-enriched regulatory sites show changes in histone acetylation and/ trimethylation abundance with respect to time of the day. To this end, we examined intensity of histone modifications in both SCN and cortex at ZT3 (day) and ZT15 (night), and compared the resulting peak profiles. We clearly observed a significant proportion of varying H3K27ac peak abundance between day and night exclusively in the SCN (Fig. S3B), with no difference in the cortex. This strongly suggests towards SCN specific *cis*-elements that confer diurnal change in chromatin accessibility, potentially to control downstream rhythmic gene transcription. This shows the importance of temporal control in genome-wide histone modifications within distinct brain regions in the context of daily timekeeping. Intriguingly, the differential intensity noted in histone modification in the SCN was only observed for H3K27ac, and not for the H3K4me3 mark, which is profoundly enriched around the gene TSS. A plausible explanation for this phenomenon could be the occurrence of H3K4me3 marks around both poised and active gene TSS (Beacon et al., 2021), whereas H3K27ac is reported to be present chiefly around active enhancers (Creyghton et al., 2010). Therefore, in our current study H3K27ac was a better marker of dynamic changes corroborating the notion that active enhancers are reliable determinants of spatiotemporally changing gene expression, which is a prerequisite for circadian timekeeping (Panigrahi & O’Malley, 2021).

To explore further the occupancy of H3K27ac-defined active enhancers in the SCN, we assessed their abundance at varied 12-hr separated time intervals (Fig. S3C), and found robust 24-hourly oscillation (Fig. 4). While this is consistent with the previously reported circadian modulation in histone modification levels observed in the peripheral clocks (Koike et al., 2012; Le Martelot et al., 2012), in our study we went a step further to assign transcripts that are under the control of these dynamic enhancers in the SCN. To address this, we executed bulk RNA-seq in parallel to histone ChIP sequencing at six distinct times of the day and noticed a high correlation between fluctuating H3K27ac levels and neighbouring rhythmic gene transcripts. On close inspection, often the phase of peak H3K27ac abundance was seen to precede the rising mRNA levels (Fig. 4f), consistent with the view that chromatin accessibility is offered by histone modification prior to achieving the required transcriptional burst (Cheng & Gerstein, 2012; Karlić et al., 2010). Furthermore, with the help of the current transcriptional dataset; we noticed that the peak of rhythmic gene expression, particularly for the core-clock genes, comes first in the SCN followed by its increase in other studied peripheral tissue(s) (Abe et al., 2022; Yamamoto et al., 2004). For example, *Per2* gene expression peaks around ZT8 (middle to late day) in the SCN whereas it is reported to attain highest level around ZT14 (early night) in the liver. Hence, the systematic SCN transcriptomics conducted in the present study reconciled with the prior studies (Hogenesch et al., 2003; Panda et al., 2002) and offered an excellent opportunity to understand the relationship between changing chromatin state and gene expression.

Given the dynamic nature of histone acetylation in the SCN, we investigated the abundance of non-coding eRNAs that can regulate target gene expression, for instance by mediating the formation of enhancer-promoter loops (Kim et al., 2015). These transcribing enhancers have also been recently discovered in multiple neuronal populations, as reported near the activity induced *Fos* gene (Carullo et al., 2020), and found to be essential for induction of the target mRNA transcription. Thus, we capitalized on the exclusive bidirectional nature of eRNA transcription (Sartorelli & Lauberth, 2020) and successfully identified a distinctive repertoire of actively transcribing eRNAs in the SCN. This was particularly challenging with regards to the size of the SCN and the limited amount of genomic material harvested from it. Nonetheless, with the high sequencing depth and biological replicates, we confidently showed the prevalence of bi-directionally transcribing non-coding eRNAs alongside H3K27ac in the SCN. Remarkably, a significant proportion of these non-coding eRNAs possessed robust daily oscillation in their expression levels, concomitant with the respective H3K27ac intensities and appeared to control the rhythmic target mRNA expression in the SCN. Our current finding is an extension to the formerly studied role of non-coding RNA (Mosig & Kojima, 2022) such as miRNA (Xue & Zhang, 2018; Zhou et al., 2021) in fine-tuning circadian clock by influencing the phase, amplitude and period of the rhythm as noted across multiple organisms. Taken together, our pioneering efforts shed light on the prevalence of cycling *cis*-regulatory elements in the master pacemaker and clearly demonstrates its role in fine-tuning gene expression machinery in the SCN.

This interwoven network of gene regulatory elements provides a hierarchical system of control over gene expression in the SCN. The dynamic change in histone acetylation followed by eRNA transcription lays the foundation for the coherent gene expression imperative for a daily timekeeping mechanism. Our present findings provide an excellent explanation how sequential layers of pre-transcriptional regulation offer timely synchronisation to the SCN transcriptional grid (Fig. 7). It also opens up the opportunity for further research on functional relationships between the molecular clock and epigenetic programming. Hence, it will be of interest to determine if genetic variation within any of these regulatory elements presents a threat to the well-orchestrated circadian timekeeping mechanism directed by the SCN. This could potentially constitute starting points to understand the association between aberrant gene-regulatory regions and the human disease pathologies such as psychiatric and neurodegenerative disorders resulting from circadian mis-alignment (Fagiani et al., 2022; Logan & McClung, 2019; Zhang et al., 2014).

## Material and Methods

### Mice

All animal studies were performed under the guidance issued by Medical Research Council in Responsibility in the Use of Animals for Medical Research (July1993) and Home Office Project Licence 19/0004. WT C57BL/6J were maintained and provided in-house by MRC Harwell. Animals were group housed (4-5 mice per age) in individually ventilated cages under 12:12hr light-dark conditions with food and water available ad libitum.

### Experimental Design

WT C57BL/6J mice (males, aged between 8 to 12 weeks) were used for SCN tissue collection as described by Jagannath et al (Jagannath et al., 2013) at 6 distinct time-points starting from ZT3 at every 4 hours, where lights on at 7 am (ZT0) and lights off at 7 pm (ZT12). We also collected cortical punches at ZT3 and ZT15 to compare and establish SCN-enriched chromatin modifications. For histone chromatin immunoprecipitation two separate biological replicates per time-point per tissue-type was collected where each biological replicate composed of 3-4 individual SCN or cortical punches. For SCN-Bulk RNA-Seq four biological replicates per time-point were collected. Likewise, each biological replicate constituted 3-4 individual SCN samples.

### Histone chromatin immunoprecipitation (ChIP) and sequencing

Chromatin immunoprecipitation (ChIP) was conducted by Diagenode ChIP-seq/ChIP-qPCR Profiling service (Diagenode Cat# G02010000). Briefly, the chromatin was prepared using the True MicroChIP Kit (Diagenode Cat# C01010130). Samples were fixed with 1% FA for 8 min. Chromatin was sheared using Bioruptor® Pico sonication device (Diagenode Cat# B01060001) combined with the Bioruptor® Water cooler for 7 cycles using a 30’’ [ON] 30’’ [OFF] settings. Then, shearing was performed in 0.65 ml Bioruptor® Pico Microtubes (Diagenode Cat# C30010011). 25μl of this chromatin was used to assess the size of the DNA fragments obtained by High Sensitivity NGS Fragment Analysis Kit (DNF-474) on a Fragment Analyzer™ (Advanced Analytical Technologies, Inc.). ChIP was performed using IP-Star® Compact Automated System (Diagenode Cat# B03000002) following the protocol of the aforementioned kit. 500 ng of chromatin were immune-precipitated using 1 μg of each of the following Diagenode antibodies H3K4me3 (C15410003; Lot Nr. A1051D), H3K27ac (C15410196; Lot Nr. A1723-0041D) and Rabbit IgG (C15410206; Lot Nr. RIG001).

Chromatin corresponding to 1% was set apart as Input. The DNA after reverse cross-linking is quantified using Qubit™ dsDNA HS Assay Kit (Thermo Fisher Scientific, Q32854). Moreover, qPCR analysis was made to check ChIP efficiency using KAPA SYBR® FAST (Sigma-Aldrich) on LightCycler® 96 System (Roche) and results were expressed as % recovery = 100*2^((Ct_input-6.64)-Ct_sample). Primers used were the following: the promoter of GAPDH (GAPDH-TSS) and myoglobin exon 2 (MBex2).

H3K4me3 and H3K27ac immune-precipitated genomic DNA along with their corresponding input samples was sent to Oxford Genomics Centre, University of Oxford for library preparation using ChIP-seq protocol and paired-end sequencing on NovaSeq6000 platform (Illumina).

### Histone ChIP –Seq mapping and peak calling

Raw sequence data in the form of a pair of fq.gz with were processed using tools on the Galaxy EU server (usegalaxy.eu) using the ChIP-Seq pipeline. Paired-end FASTQ files were quality assessed by removing low quality bases (Phred<20) and trimmed using FastQc and Trimmomatic (v0.36), respectively. FASTQ files containing trimmed sequences were then aligned to mm10 genome assembly to generate binary alignment map (BAM) files with Bowtie2 (v2.3.4.1). Aligned files were filtered for minimum mapping quality (MAPQ)>20 by SAMtools (v1.8), and used for peak calling by MACS2(v 2.1.1.20160309) with the options – qvalue 0.05 –gsize mm:1.87e9 –format BAMPE. Finally, peaks for each brain region and histone modification (H3K4me3, H3K27ac) were analysed for differential binding by a Bioconductor package Diffbind (v2.10.0).

### Bulk RNA-sequencing

Bulk RNA-sequencing (RNA-seq) was carried out at Oxford Genomics Centre, University of Oxford. RNA was extracted, DNase treated and purified (RNeasy, Qiagen) for four biological replicates per time-point. 500ng-1ug of total RNA underwent quality control (Nanodrop) and libraries were prepared for directional Ribodepleted-RNA Sequencing using NEBNext reagents. RNA-Seq libraries underwent sequencing (150 bp paired-end directional reads: ~ 50 million reads/sample) on NovaSeq6000 (Illumina) platform.

### Bulk RNA-seq data analysis

Paired-end FASTQ files were quality assessed (Phred<20 removed) with FastQC and Illumina adapters were trimmed with TrimGalore (v0.4.3). Then, the reads were aligned to mm10 genome assembly using STAR (v2.7.8a) with MAPQ value for unique mappers set to 60. Binary alignment map (BAM) files were used to generate read counts per gene by FeatureCounts via Samtools (v1.11). Finally, limma-voom method (Liu et al., 2015) from the Bioconductor package-limma (v3.48.0) was adopted to quantify differential gene expression and normalised logarithmic CPM values were generated for downstream analysis.

### eRNA identification

Enhancer identification was performed by investigating H3K27ac sites as described above. MACS2 –defined and Diffbind identified differential H3K27ac peaks coupled with total RNA-Seq BAM files were used to predict bidirectional eRNA transcription using Seqmonk software (Babraham Institute) as previously reported (Carullo et al., 2020). Briefly, for eRNA identification we focussed on the intergenic H3K27ac enrichment sites as intragenic sites posed risk of alternate promoters and other regulatory elements plus nascent gene transcription. Any differential intergenic H3K27ac site that showed bidirectional transcription was further assessed for eRNA-target gene pairs. These were identified by mapping gene promoters within 2 Mb upstream or downstream from the centre of the peak. CPKM values for each identified eRNA and associated gene were correlated using Pearson’s correlation in R. eRNA-gene pairs with correlations > 0.5 were considered as high confidence pairs.

### HOMER motif analysis

Differential region or time-point specific genomic positions were compiled into BED files. The findMotifsGenome.pl function within HOMER (v4.11) package was used to identify enriched motifs and their corresponding transcription factors with options size 1000 –len 8,10,12 –mask –preparse –dumpfasta. For interested region specific motifs such as SCN, genomic bed files from the cortical region was used as the background and vice-versa.

### Gene annotation

Gene list derived from bioconductor based ChIPseeker (v1.28.3) was fed into Database for Annotation, Visualization and Integrated Discovery (DAVID) tool (Dennis et al., 2003). The functional annotation chart based on KEGG pathway and Gene Ontology (GO∷BP) was plotted with the help of ggplot 2 package in R(4.0.5).

### Analysis of oscillating H3K27ac signal, gene and eRNAs

Logarithmic counts per million (CPM) values across all time-points for H3K27ac occupancy, gene and eRNA feature were analysed for significant circadian oscillations using JTK_CYCLE (Wu et al., 2016) and ECHO (De Los Santos et al., 2020). For H3K27ac oscillation, significantly differential H3K27ac site between any two anti-phasic (12 hour apart) were compiled using Bedtools(v2.30.0). Normalised CPM values from compiled 1021 intervals was used for 24 hour rhythmicity assessment.

### RNA extraction and RT-qPCR

Total RNA was extracted from the SCN tissue as described above at 3 distinct time-points; ZT3, ZT15, ZT23. For each time-point RNA was extracted from 2-4 biological replicates, where each biological replicate comprises of 3-4 individual SCN punches. 100 ng RNA per sample replicate was reverse-transcribed (iScript cDNA Synthesis Kit,Bio-Rad; 5 min at 25°C, 20 min at 46°C, 1 min at 95°C, hold at 4°C). The synthesized cDNA was used for RT_qPCR using Applied Biosystems 7500 Real-time PCR instrument (SsoAdvanced Universal SYBR Green Supermix, Bio-Rad; 1 ul cDNA in 20 ul, 30 sec at 95°C, 40 × 15 sec at 95°C and 60 sec at 60°C, followed by melt curve analysis 65-95°C in 0.5°C increments at 5 sec per step) in triplicates for genes of interest (*Vipr1*, eRNA (chr9: 123334375-123335717)). 7500 Applied Biosystems Software (v2.3) was used to obtain the relative quantification values and examine melt curves. All RT-qPCR data was normalized to *Rpl-32* reference control and analysed using standard curve method (Larionov et al., 2005) The primers used for *Vipr1*: TCAACAACGGGGAGACAGAC (Forward) and GGCCATGACGCAATACTGGA (Reverse), eRNA: TGGTTAAGAGCGCCTACAGC (Forward) and GCTGTCTTCAGACACTCCAGA (Reverse), *Rpl-32*: AGGCACCAGTCAGACCGATA (Forward) and TGTTGGGCATCAGGATCTGG (Reverse), were designed and screened for target specificity with the National Center for Biotechnology Information’s (NCBI) Basic Local Alignment Search Tool (BLAST). The primer sets were validated prior to use (requirements: no predicted off-targets, primer efficiency 85120%, no significant signal in non-template control, single peak in melt curve, and single band at the predicted size when separated via agarose gel electrophoresis).

### Statistical Analysis

Gene expression differences from qPCR were plotted and analysed with Graphpad Software (Prism). Pearson correlation between target mRNA and eRNA across the tested time-points, set at alpha =0.05.

## Data Availability

The data discussed in this publication have been deposited in NCBI’s Gene Expression Omnibus (Edgar et al., 2002) and are accessible through GEO Series accession number GSE217943 (https://www.ncbi.nlm.nih.gov/geo/query/acc.cgi?acc=GSE217943).

## Author contribution

Conceptualization: AB, GB, PMN

Methodology: AB, GB

Investigation: AB, PMN

Formal analysis: AB

Visualization: AB, GB, MHH, PMN

Funding acquisition: PMN

Supervision: GB, MHH, PMN

Writing – original draft: AB, PMN

Writing – review & editing: AB, GB, MHH, PMN

## Acknowledgements

We thank the staff of the Mary Lyon Centre and core services at MRC Harwell Institute for assistance with mouse studies. We thank Professor Felix Naef, EPFL, Lausanne for his valuable suggestions and timely inputs. We also thank Dr Michelle Simon and Richard Reeves for data accessibility on UCSC genome browser. This research was supported by the Medical Research Council (MC_U142684173) grant to PMN, the Hastings lab was supported by Medical Research Council (MC_U105170643).

## Supplementary Materials

Supplementary Figures: Fig. S1 to S6

Supplementary data: Table S1 to S13.

## Supplementary Figures

**Fig S1:**
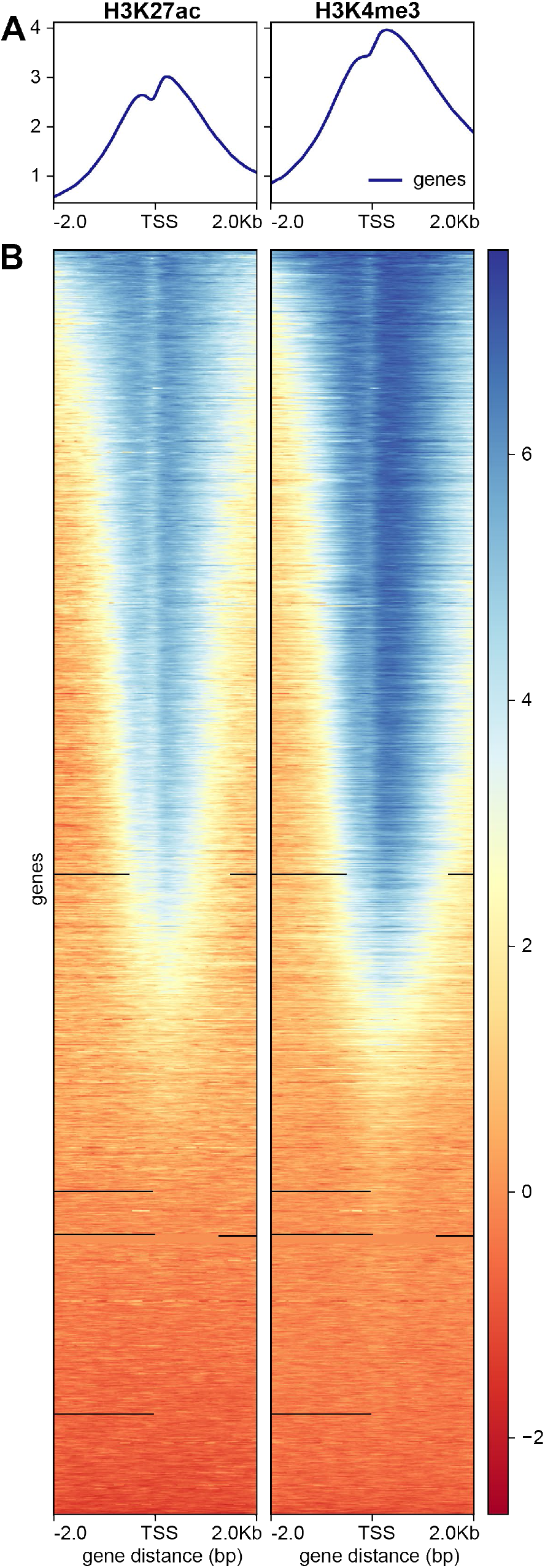
Distribution of H3K4me3 and H3K27ac sites around TSS. (A) Plot profile and (B) Heatmap showing distribution of H3K4me3 and H3K27ac sites up to 2Kb upstream and downstream from transcription start site (TSS).

**Fig. S2.**
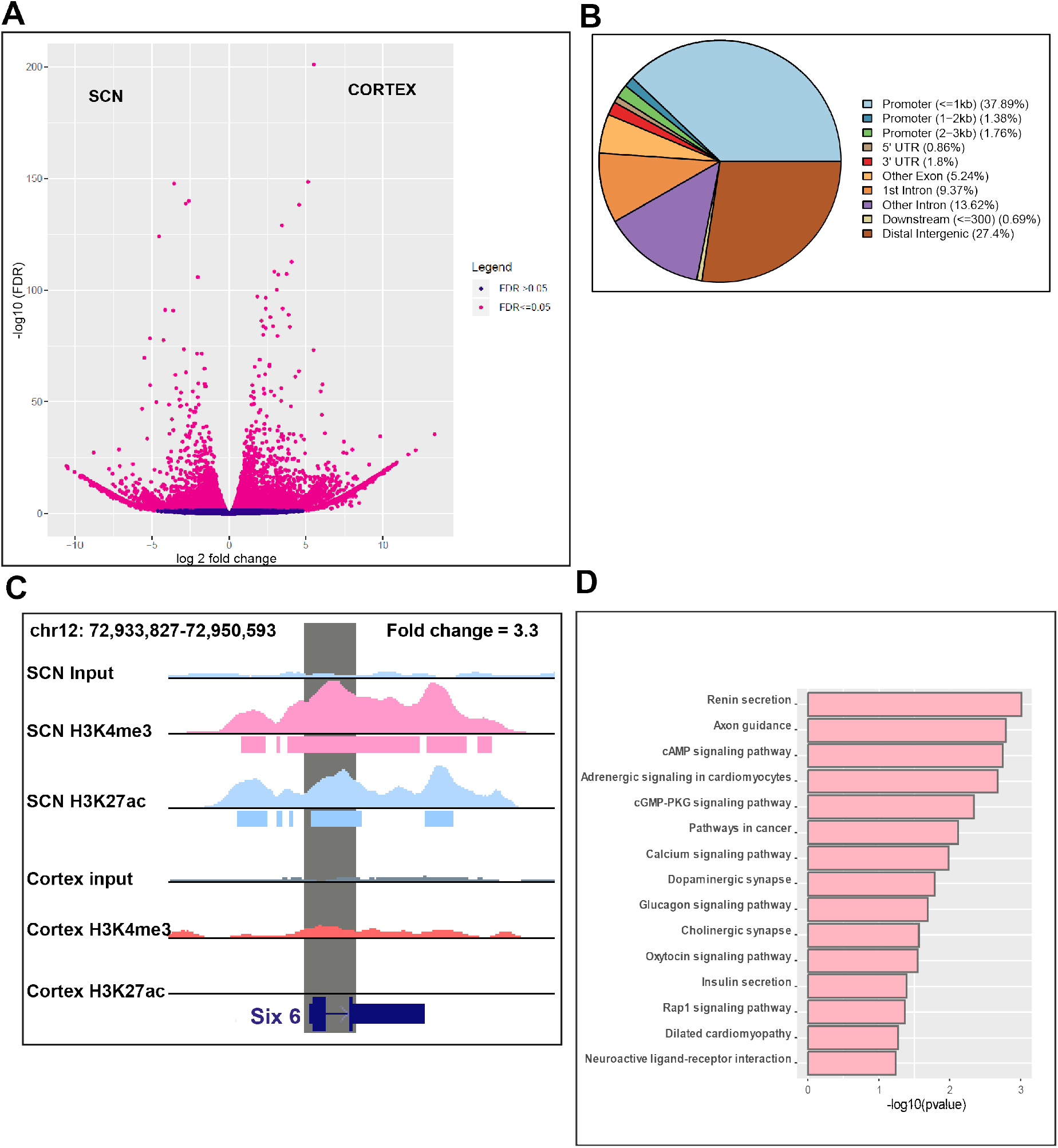
Differential H3K4me3 sites between SCN and Cortex. (A)Volcano plot showing fold change and false discovery rate (FDR) for differential H3K4me3 sites between SCN and Cortex as computed by Diffbind. (B) Genomic feature distribution of differential H3K4me3 sites between SCN and Cortex (n = 10577).(C) UCSC genome browser tracks showing H3K4me3 and H3K27ac normalised ChIP-seq coverage and peaks around *Six6* locus (shaded grey), demonstrating H3K4me3 enrichment in SCN (pink Vs red genome coverage). (D) Functional annotation of nearest neighbouring gene (TSS) to SCN enriched H3K4me3 sites (fold change > 5) using KEGG pathway (DAVID)

**Fig. S3.**
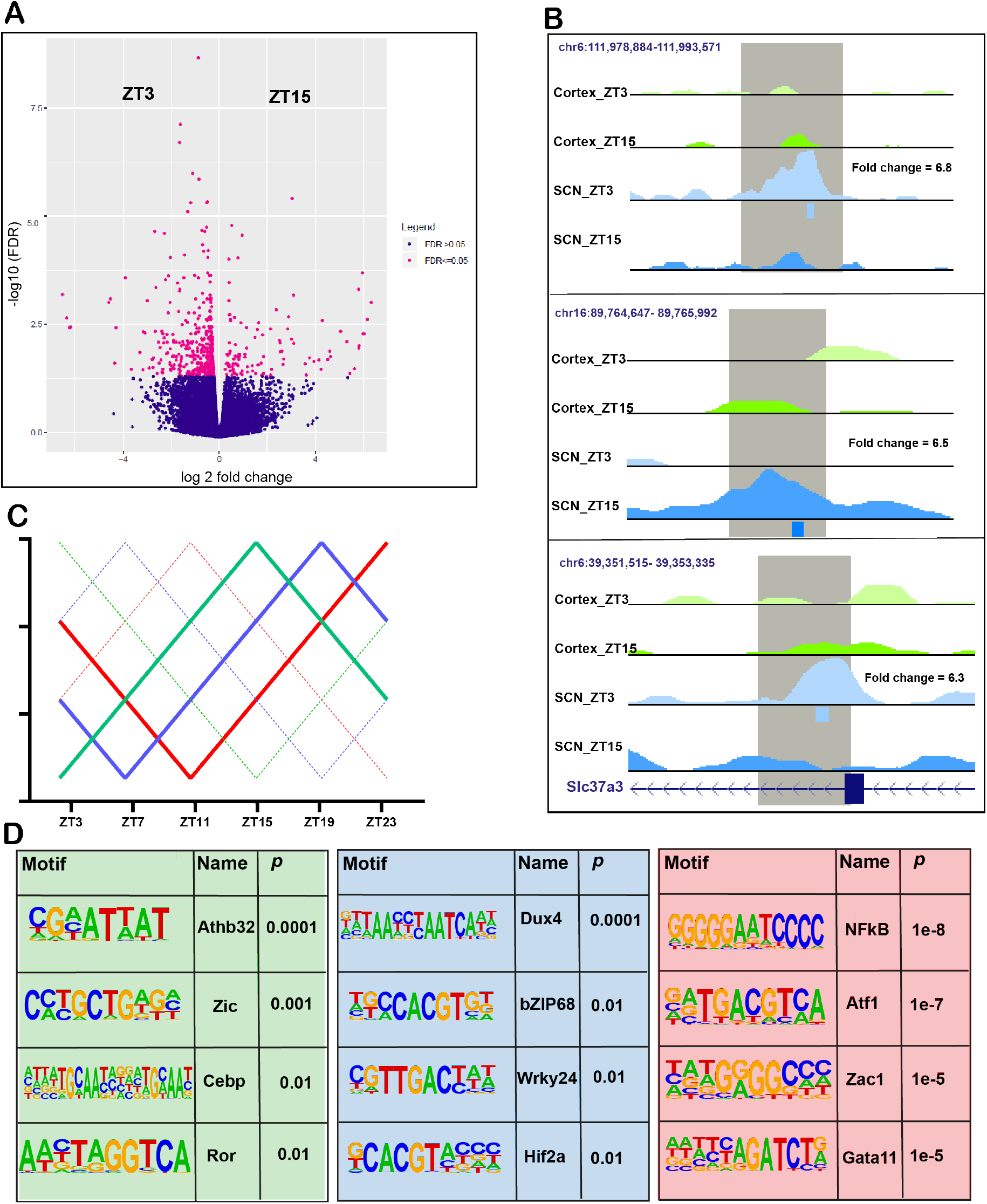
Differing H3K27ac occupancy at anti-phasic timepoints. (A)Volcano plot showing fold change and false discovery rate (FDR) for differential H3K4me3 sites between ZT3 and ZT15 (n = 551) in Cortex. (B) UCSC genome browser images of three separate genomic regions showing differential ZT3 Vs ZT15 H3K27ac abundance specifically in SCN, fold change and chromosome positions indicated per region. (C) Representative model illustrating anti-phasic timepoint comparisons with bold and dashed lines for ZT3 Vs ZT15 (green), ZT7 Vs ZT19 (blue) and ZT11 Vs ZT23 (red). (D) Enriched motif at differential H3K27ac sites observed between ZT3 Vs ZT15 (green shaded), ZT7 Vs ZT19 (blue shaded) and ZT11 Vs ZT23 (red shaded).

**Fig. S4.**
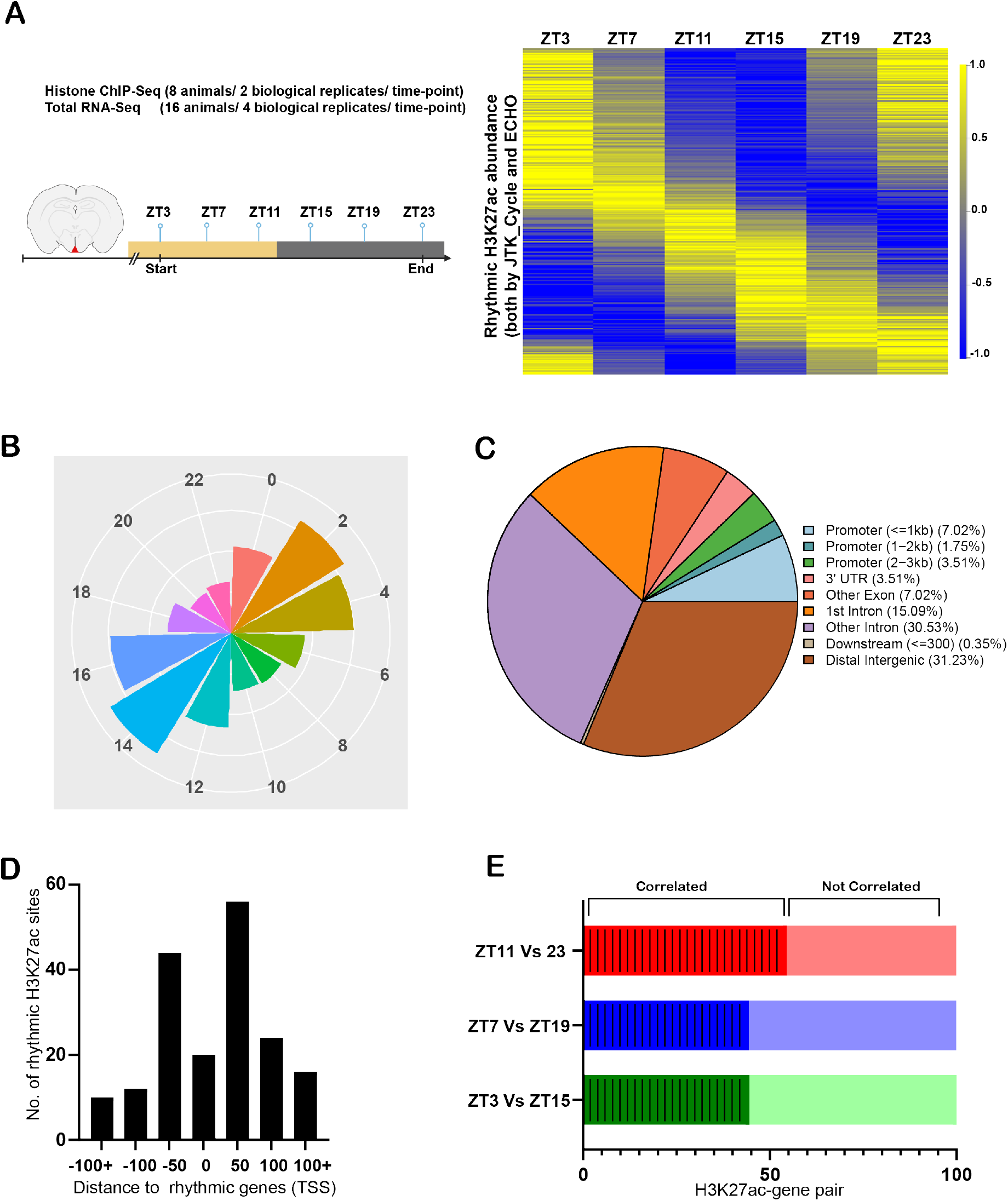
Diurnal variation in H3K27ac occupancy. (A) Illustration of SCN tissue collection for histone ChIP-seq and RNA-seq at six distinct time-points and heatmap showing rhythmic (both ECHO and JTK_CYCLE) abundance of H3K27ac across the day. (B) Rose plot showing phase (peak expression) distribution (12 groups) of rhythmic genes in SCN. The area of each wedge is proportional to number of rhythmic genes in that group. (C) Genomic feature distribution of rhythmic H3K27ac intervals as analysed by ChIPseeker. (D) Bar plot indicating the number of rhythmic H3K27ac sites and their distance from closest rhythmic gene/TSS (Kb). (E) Proportion of correlated (Pearson correlation = 1) and not correlated (Pearson correlation = −1) H3K27ac-gene levels for each compared antiphasic time-points.

**Fig. S5.**
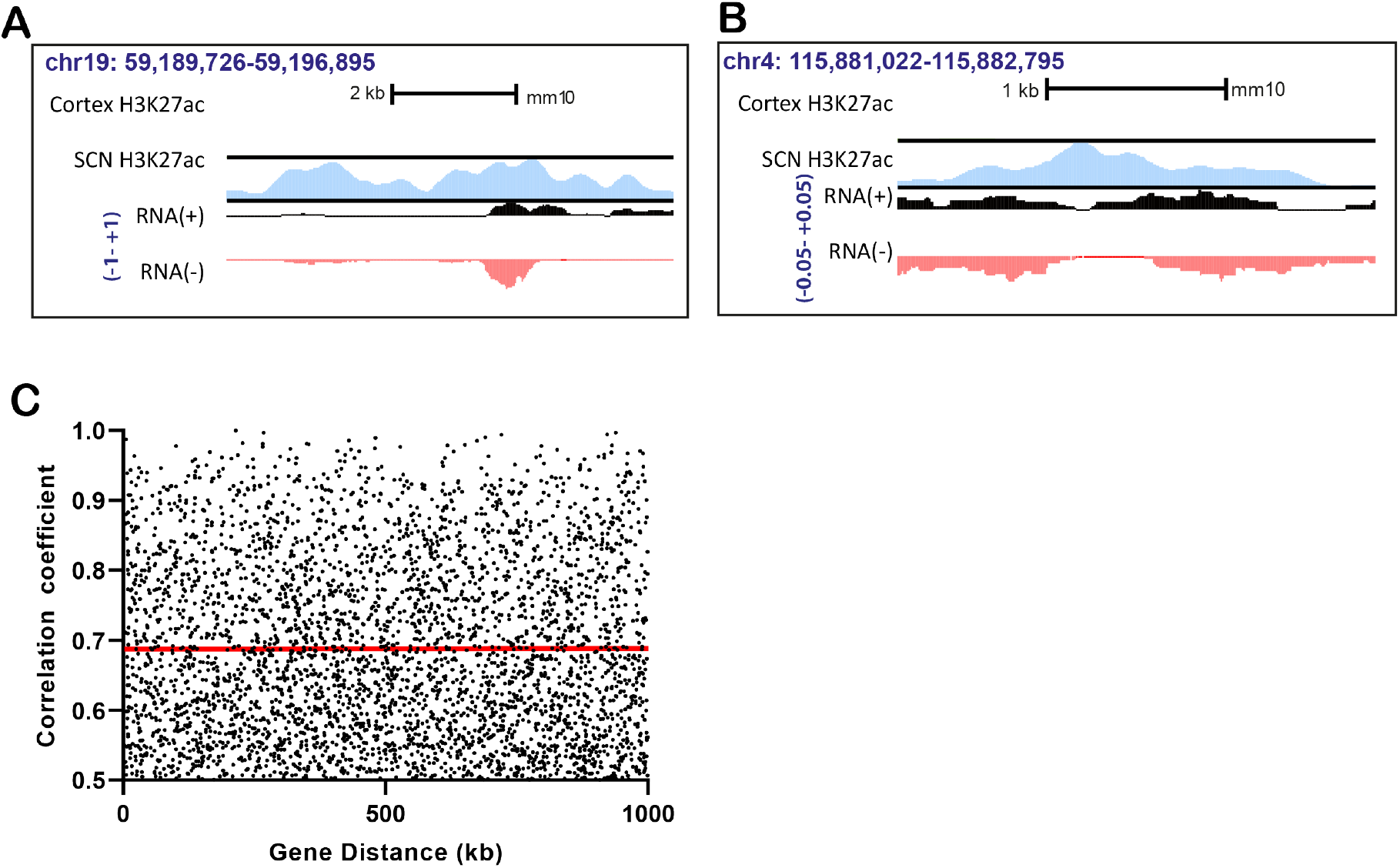
Enhancer RNA identification in SCN. (A and B) Genome browser images showing bidirectional transcription (signal as normalised CPM value from + (black) and − (red) RNA strands) arising from H3K27ac marked SCN enhancer peak (blue), at marked mm10 chromosome coordinates. (C) Scatter plot showing no relation between gene distance and strength of correlation (*r* > 0.5) as indicated by red line.

**Fig. S6.**
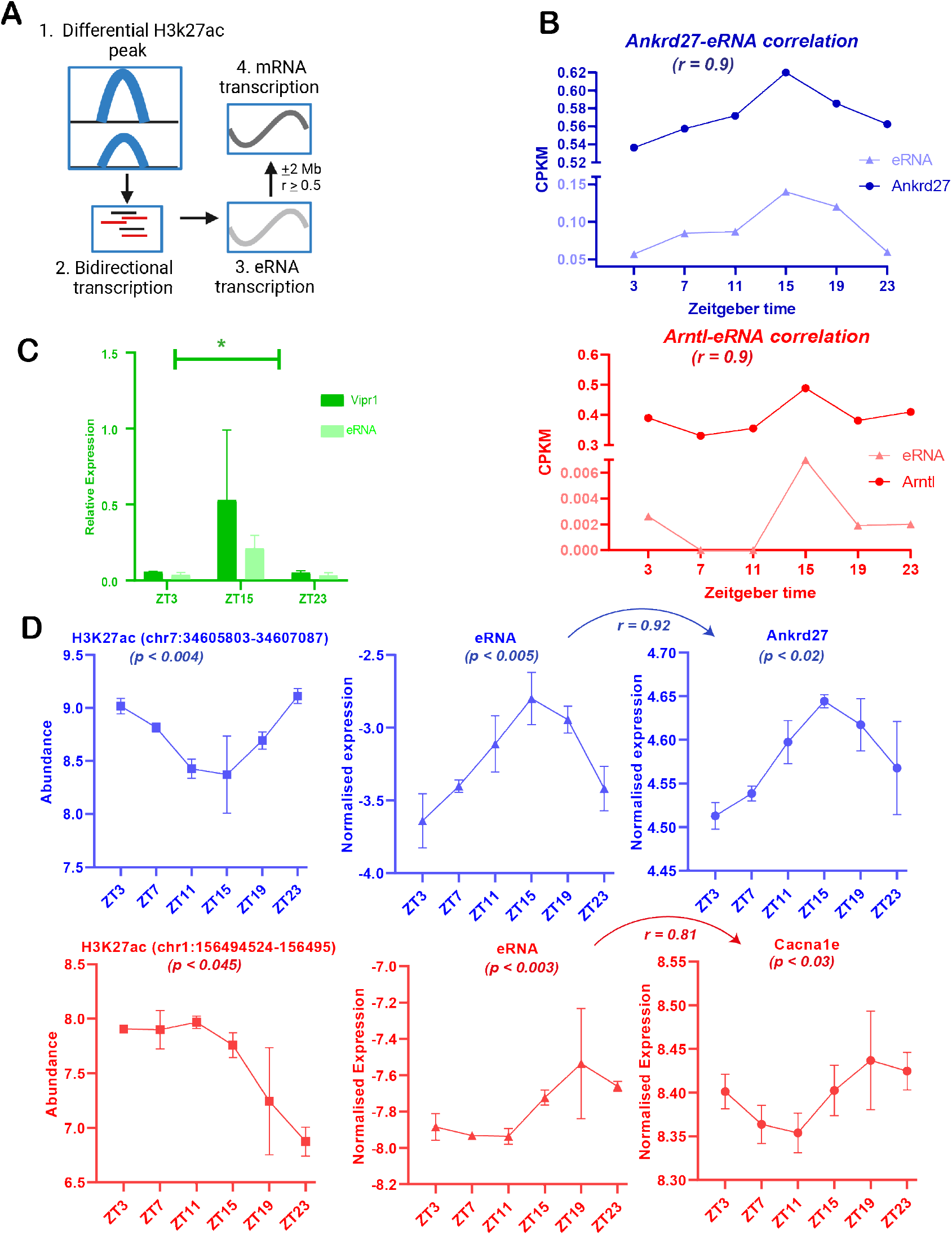
Differential H3K27ac abundance linked with eRNA and target mRNA transcription. (A)Schematic of eRNA identification from differential H3K27ac peaks (B) eRNA and positively correlated *(r* > 0.5) target mRNA expression (CPKM) during the day. (C) Relative expression of eRNA (chr9: 123334375-123335717) and *Viprl* mRNA analyzed by qPCR at ZT3, 15 and 23, showed perfect correlation in peak and trough expression levels (Pearson *r* = 0.99, p < 0.01 as denoted by *). (D) Representative examples of differential H3K27ac sites (chromosome coordinates indicated at top) observed across ZT7 Vs ZT19 (blue) and ZTI I Vs ZT23 (red), showing rhythmic H3K27ac abundance along with cyclic eRNA and target gene expression (log CPM) as analysed by ECHO (p<0.05) with indicated Pearson correlation coeeficient (*r*).

